# Development of a novel epigenetic clock resistant to changes in immune cell composition

**DOI:** 10.1101/2023.03.01.530561

**Authors:** Alan Tomusiak, Ariel Floro, Ritesh Tiwari, Rebeccah Riley, Hiroyuki Matsui, Nicolas Andrews, Herbert G. Kasler, Eric Verdin

**Author notes:** Contributing authors.

## Abstract

Epigenetic clocks are age predictors that use machine-learning models trained on DNA CpG methylation values to predict chronological or biological age. Increases in predicted epigenetic age relative to chronological age (epigenetic age acceleration) are connected to aging-associated pathologies, and changes in epigenetic age are linked to canonical aging hallmarks. However, epigenetic clocks rely on training data from bulk tissues whose cellular composition changes with age. We found that human naive CD8^+^ T cells, which decrease during aging, exhibit an epigenetic age 15–20 years younger than effector memory CD8^+^ T cells from the same individual. Importantly, homogenous naive T cells isolated from individuals of different ages show a progressive increase in epigenetic age, indicating that current epigenetic clocks measure two independent variables, aging and immune cell composition. To isolate the age-associated cell intrinsic changes, we created a new clock, the IntrinClock, that did not change among 10 immune cell types tested. IntrinClock showed a robust predicted epigenetic age increase in a model of replicative senescence *in vitro* and age reversal during OSKM-mediated reprogramming.

## Introduction

Epigenetic clocks, age predictors based on DNA methylation levels at selected CpG loci, have grown in popularity as a tool to study aging and predict health outcomes in humans. The first epigenetic clocks developed by Hannum et al.^1^ and Horvath^2^ showed remarkably high accuracy (R > .90) in predictions of chronological age. These “first-generation” epigenetic clocks provide unique biological insights into the aging process. For example, some but not all forms of senescence accelerate epigenetic clock age predictions^3^. Using a later clock trained on chronological age, Kabacik et al.^4^ identified nutrient sensing, mitochondrial activity and stem cell composition as being associated with epigenetic aging but not telomere attrition or genomic instability. A recent report demonstrated the development of an epigenetic clock effective at predicting age across a variety of species, providing evidence for a shared mammalian aging program^5^.

More recently, “second-generation” clocks designed to predict phenotypic aging measures have been developed. These clocks, including PhenoAge^6^ and GrimAge^7^, show strong associations with diseases, such as depression^8^ and mortality^9^. DunedinPACE is a similar marker of phenotypic aging that captures the pace of aging rather than the accumulation of aging^10^. These clocks show promise as markers of physiological aging, but their two-step construction methodology (training a DNA methylation predictor on measures of phenotypic rather than chronological age) adds a secondary layer of complexity to interpretation.

Given the ability of epigenetic clocks to detect aging phenotypes across species and levels of organization that include cells, tissues, and organs, there is significant interest in understanding the underlying mechanism(s) enabling their function. Recent preprints have been released on this topic, notably including one by Levine et al.^11^ that suggests epigenetic clocks are composites of different modules characterized by their changes during aging and reprogramming. Novel epigenetic clocks have been developed that seek to capture the aging phenomenon in more defined ways, including by identifying CpG sites predicted to be causal by Mendelian randomization^12^ or those capturing purely stochastic variation^13^. These clocks are informative about aspects of the aging process and have the potential to be particularly well-suited for certain use cases.

One major challenge in understanding the mechanism(s) underlying epigenetic clocks is the confounding effect of age-related changes in cell-type composition of many tissues. While changes in cell-type composition are an important part of aging, they can make interpreting epigenetic clocks more difficult as the relevant CpG sites may be cell-type-specific markers rather than those affecting cell-intrinsic aging. Most epigenetic clocks are trained largely on blood, which sees a drop in naïve CD8^+^ T cells with age and a corresponding increase in more terminally differentiated memory T-cell types^14^. Some clocks may be more impacted by changes in cell-type composition than others, depending on how they were constructed^15^. T-cell and NK (natural killer) cell activation have been implicated as major drivers in epigenetic clock progression^16^.

Other approaches have been explored to create epigenetic age predictions that are less sensitive to changes in cell type composition. Most notably, residuals from regression models that include epigenetic age and proportions of several blood cell types have been used to generate an “ intrinsic epigenetic age acceleration” measure^17^. While the resulting measure is cell-type independent, it becomes challenging to biologically interpret as the underlying signal is derived from a mixture of CpG sites that can be either cell type-independent or cell type-dependent. Other modern approaches include the development of single-cell epigenetic clocks^18,19^, though the underlying technology will require further maturing before it can match the sensitivity and accuracy of bulk measurement-based clocks.

In this work, we report our analysis of the differences in epigenetic age predictions derived from four epigenetic clocks (Hannum^1^, Horvath^2^, Horvath Skin and Blood^20^, and PhenoAge^6^) for cytotoxic CD8^+^ T cells at different stages of differentiation. We found that human naïve CD8^+^ T cells, which decrease in humans during aging, exhibit an epigenetic age 15–20 years younger than effector memory CD8^+^ T cells isolated from the same individual. Interestingly, naïve T cells isolated from individuals of different ages still show a progressive increase in epigenetic age. Based on these observations, which indicate, as predicted, that current epigenetic clocks measure two independent variables, aging and immune cell composition, we created a new clock, the IntrinClock, that does not change among 10 immune cell types tested. Remarkably, this clock shows an increase in a model of replicative senescence *in vitro* and shows decreased aging during OSKM reprogramming. Lastly, we investigate the IntrinClock’s applicability for use in studying and detecting the effects of cell-intrinsic perturbations on aging.

## Results

### Existing epigenetic clock age predictions depend on CD8^+^ T-cell differentiation state

In humans, CD8^+^ T cells decrease in frequency, with a particularly pronounced loss of naive T cells during aging^21^. We used a negative bead-based selection method to isolate total T cells from seven donors of varying ages, all of whom were positive for cytomegalovirus (CMV^+^). We then used FACS to isolate CD8^+^ naive (CD8^+^ CD28^+^ CD45RO^-^), CD8^+^ central memory (CD8^+^ CD28^+^ CD45RO^+^), CD8^+^ effector memory (CD8^+^ CD28^-^ CD45RO^+^), and CD8^+^ terminal effector memory RA^+^ (CD8^+^ CD28^-^ CD45RO^-^) cells (Figure 1A). After DNA isolation and profiling using the Illumina Infinium MethylationEPIC™ platform, we noted a distinct clustering of CD8^+^ naive cells away from CD8^+^ central memory (CM), effector memory (EM), and terminal effector memory RA^+^ cells (TEMRA) (Figure 1B) in UMAP analysis. Horvath clock epigenetic ages were measured in each of the CD8 T-cell subsets and found to correlate with age across every subset. However, strikingly, naive T cells consistently showed a significantly younger epigenetic age than other CD8^+^ subsets (Figure 1C). This result suggests that epigenetic clock measurements are affected by CD8^+^ T-cell differentiation. Equally interestingly, naive CD8^+^ T cells from individuals of different chronological age showed an increase in epigenetic age that was parallel to chronological age but consistently lower than the chronological age (Figure 1C). The same observation was made for CMs, EMs and TEMRAs except that these cells’ epigenetic age appeared closer to the chronological age of the donors.

**Figure 1.**
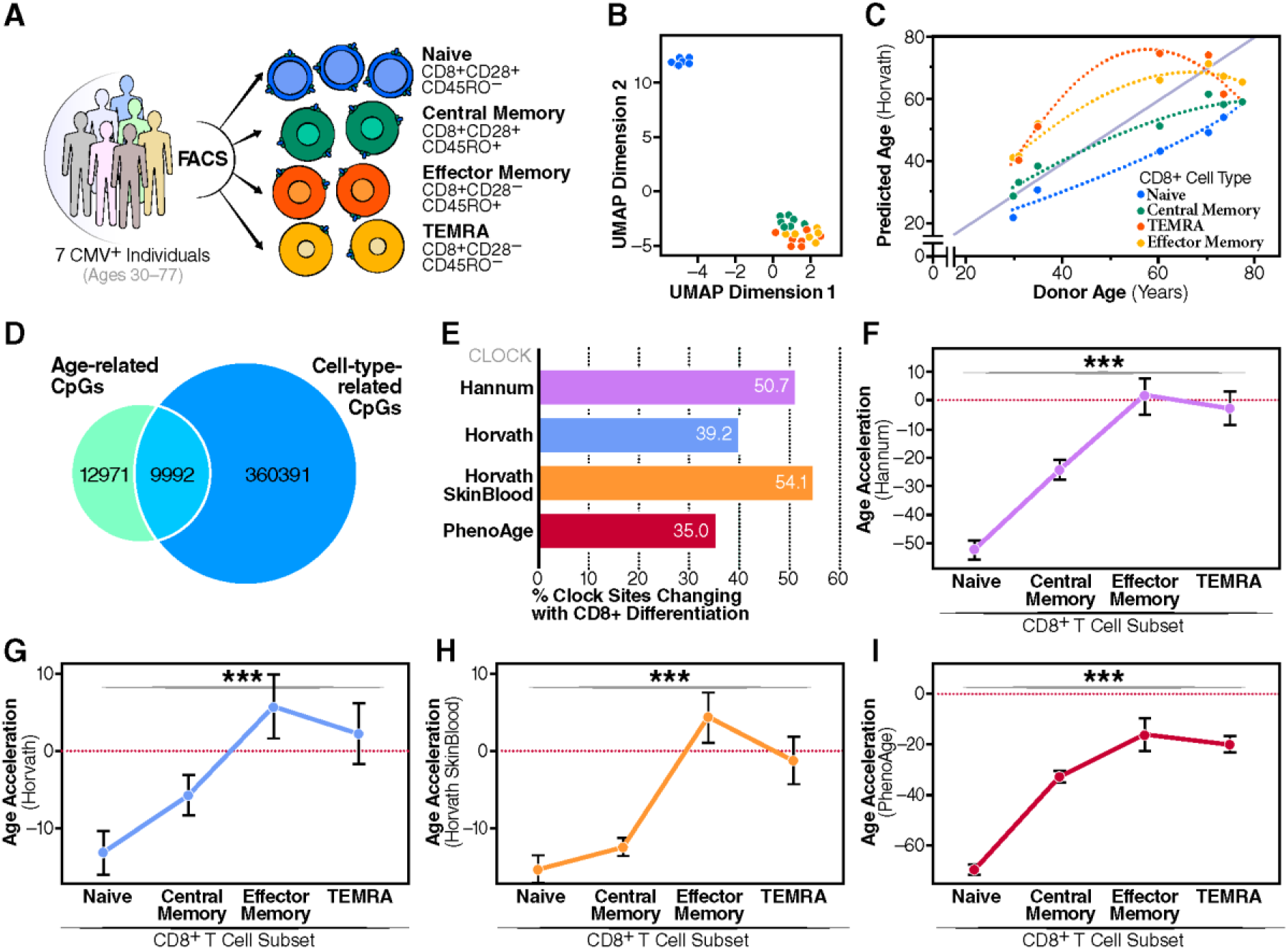
CpG site changes during T-cell differentiation. **a**, Experimental design for determining impact of CD8^+^ differentiation on epigenetic clock age prediction. **b**, UMAP dimensionality reduction of CD8^+^ DNA methylation profiles. **c**, Differences between predicted epigenetic age as a function of donor age and CD8^+^ T-cell subset. **d**, Comparison of shared CpG site changes between age in CD8^+^ T cells and CD8^+^ cell subset. **e**, Percent of sites in four epigenetic clocks that are altered by CD8^+^ T-cell differentiation. **f-i**, Comparison of the (f) Hannum, (g) Horvath, (h) Horvath skin and blood, and (i) PhenoAge epigenetic age acceleration predictions for four CD8^+^ T-cell subsets. *** ANOVA p-value less than .001.

Next, using differential methylation analysis on methylation M-values, we identified 22,963 CpGs that changed with age and 370,383 CpGs that changed between naive CD8^+^ T cells and CD8^+^ CM, CD8^+^ EM, or CD8^+^ TEMRA cells. Of the 22,963 aging-related CpGs, 9,992 were also affected by differentiation (Figure 1D). To understand how this could affect epigenetic clock predictions, we investigated the proportion of CpG sites used for epigenetic age prediction in the Hannum, Horvath, Horvath Skin and Blood, and PhenoAge clocks that we identified were affected by CD8^+^ T-cell differentiation. In all four clocks, more than a third of the predictive sites were changed with differentiation (Figure 1E), and all four had a difference in age acceleration for CD8^+^ T-cell subsets. In all clocks, CD8^+^ TEMRA and CD8^+^ EM cells were predicted to be older than CD8^+^ CM cells, which were predicted to be older than CD8^+^ naive cells (Figures 1F-1I). The differences in epigenetic ages among the CD8^+^ T-cell subsets varied among clocks. For example, PhenoAge predicts CD8^+^ naive cells to be over 60 years younger than the donor chronological age, but the difference was much smaller for both Horvath clocks with an epigenetic age prediction of only approximately 12 years lower than chronological age (Figures 1F - 1I).

### Development of a novel epigenetic clock (IntrinClock) resistant to changes in immune cell composition

Given the overlap of DNA methylation signatures of cellular aging and CD8^+^ differentiation, we sought to create a new epigenetic clock that is unaffected by changes in immune cell composition. We began by generating a database of 14,601 DNA methylation samples from 71 different datasets^1,22–90^, generated on either the Illumina Infinium™ HumanMethylation450 (450K) or the Illumina Infinium™ MethylationEPIC (EPIC) array, all sourced from the Gene Expression Omnibus (GEO) database or the Genotype-Tissue Expression project (GTEx) (Supplementary Table 1). The number of samples per dataset ranged from six to 1,218, with a mean number of samples per dataset of 213 (Figure 4A). The distribution of sexes was approximately equal (Figure S4B). Samples were derived from a variety of tissues with the majority from blood (Figure S4C), and the DNA methylation assay platform was split roughly evenly between the 450K and the EPIC array. (Figure S4D).

Once the database of samples was assembled, we performed a series of filtering and quality control steps. We filtered out all samples that were missing more than 10% of CpG sites measured by the 450K array, those that were derived from cancerous tissue, and those that were derived from germline tissues. We then removed outliers, defining outliers as those with principal components more than two interquartile ranges away from the mean (Figure 2B). After performing a random 75-25 training/test split, 9104 samples were used to train the model and 2994 were used to validate it.

**Figure 2.**
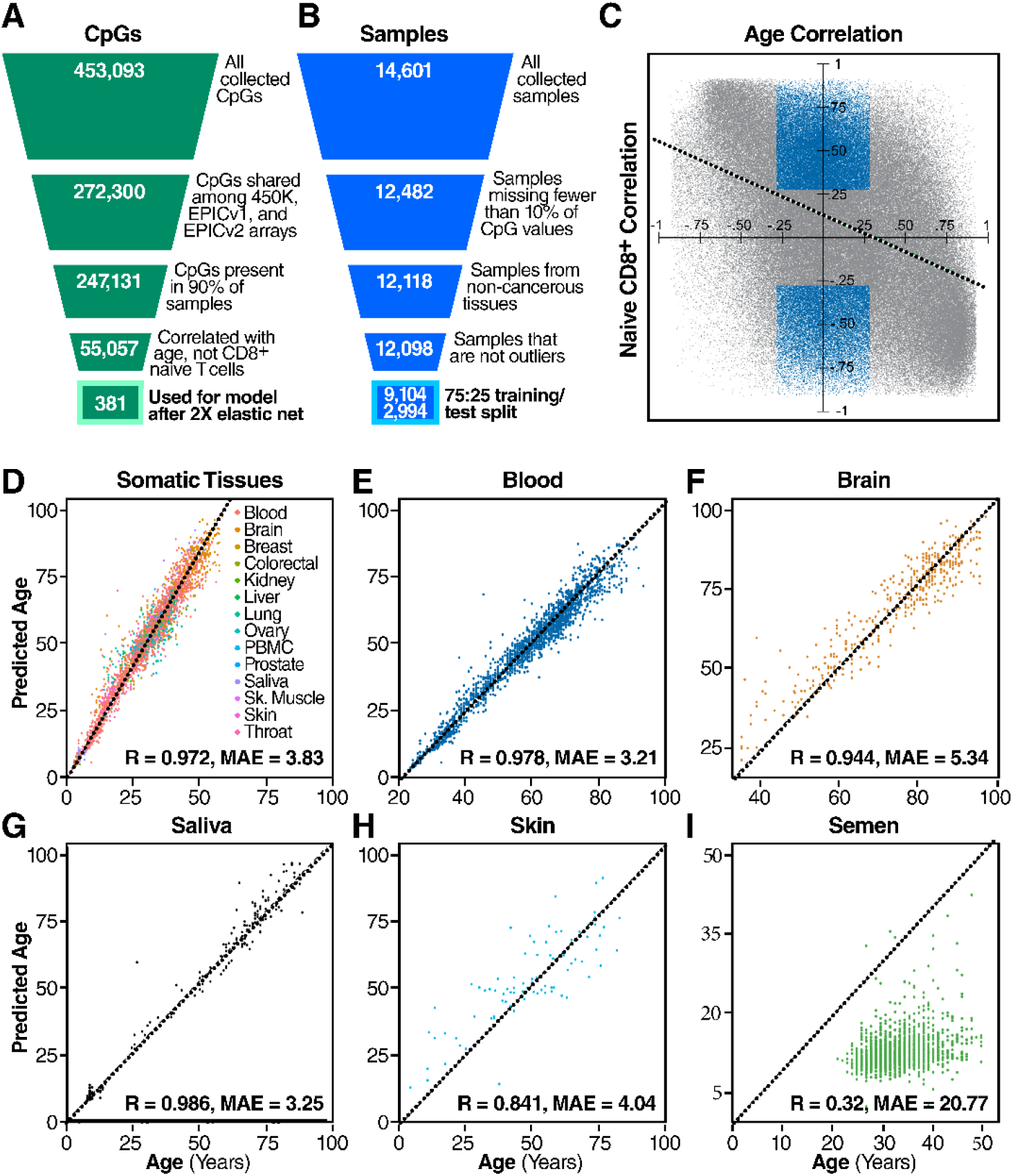
Preparation of tissues. **a**, Filtering strategy for CpG sites. **b**, Filtering strategy for samples. **c**, Visualization of the filtering process for differentiation-independent age-related CpGs. Blue CpGs (those correlated with age but not with being a naive cell) were included in the feature set, whereas gray CpGs were not. Green dashed line indicates linear least-squared regression line of relationship between CpG age correlation and CpG CD8^+^ naive cell correlation. **d**, Correlation between age and IntrinClock predicted age in a variety of tissues from the test set. **e-h**, Individual correlation plots for specific tissues in the test set. **i**, Epigenetic age vs. chronological age correlation plot for semen samples.

Given the unique methylation pattern (Figure 1B) and quiescent biology^91^ of naive CD8^+^ T cells, we aimed to use them as a basis on which to eliminate CpGs linked to CD8^+^ T-cell differentiation and performed additional filtering steps. When constructing our database of DNA methylation data, we initially collected all CpG sites measured by the 450K array for all samples. To increase reliability, we first filtered out CpG sites that were present in fewer than 90 percent of samples. To ensure forward compatibility, we also included only CpG sites that were present on the Illumina Infinium™ MethylationEPICv2.0 array. Next, we opted to remove any CpG sites that were correlated with a sample being a naive CD8^+^ sample (R > .3) within our CD8^+^ subset data (i.e., CpG sites whose methylation patterns were distinct in CD8^+^ naive cells as compared to CD8^+^ CM/EM/TEMRA cells). We also opted to include only those CpG sites correlated with age (R > .3) (Figure 2C), to decrease the search space for the elastic net algorithm to identify age-predictive sites. Interestingly, we observed a negative correlation-of-correlations between the age correlation and naive CD8^+^ correlation of CpG sites (R = - .45) (Figure 2C), indicating that CpG sites that are hypermethylated with age tend to be hypomethylated in naive CD8^+^ cells, and vice-versa. We utilized the elastic net algorithm on the remaining 55,896 CpGs to generate a new epigenetic clock based on 410 CpG sites. To increase accuracy and reduce the number of necessary prediction sites, we used a novel approach whereby we employed the elastic net algorithm a second time on the training data filtered only on the 410 CpG sites used for the clock. This reduced the number of predictive CpG sites in the final model (*IntrinClock*) to 381, and reduced error by approximately 3 months (Figure S5).

### IntrinClock is accurate across tissues, and its age predictions are not affected by adaptive immune cell compositional changes

Next, we tested the IntrinClock on a variety of tissues in the test set and observed high overall prediction accuracy (R ∼ .972, mean absolute error (MAE) ∼ 3.83) (Figure 2D). Age prediction errors on blood and saliva were particularly low (MAE ∼ 3.25, MAE ∼ 3.21, respectively) (Figure 2E, 2G). Tissues with less immune infiltration also had high epigenetic age correlations with chronological age (R ∼ .944 for brain, R ∼ .841 for skin). We were interested in discovering whether the IntrinClock would predict chronological age in semen samples, as previous epigenetic clocks have shown significant age deceleration in sperm^2^. We found that epigenetic age predictions of semen had only a weak correlation with chronological age (R ∼ .32), and the predicted age of sperm samples, using a previously generated dataset^92^, appears to consistently be ∼ 12 (Figure 2I).

Importantly and as expected, IntrinClock applied to our generated CD8^+^ DNA methylation data showed no epigenetic age prediction differences among CD8^+^ T-cell subsets (Figure 3A). As these samples were included in the training set for clock construction, we validated our approach on two external datasets^93,94^ with CD8^+^ naive and CD8^+^ EM DNA methylation data and found no differences in epigenetic age (paired t-test p-value > .05) (Figure 3B). We also tested whether our clock could find a shift in epigenetic age between CD4^+^ naive and CD4^+^ CM cells, as the proportion of CD4^+^ naive cells also decreases with age^95^. Using two external data sets^96,97^, we discovered no evidence for a shift in epigenetic age between CD4^+^ naive and CM cells (Figure 3C) (paired t-test p-value > .05), despite our filtering strategy being based only on CD8^+^ cells.

**Figure 3.**
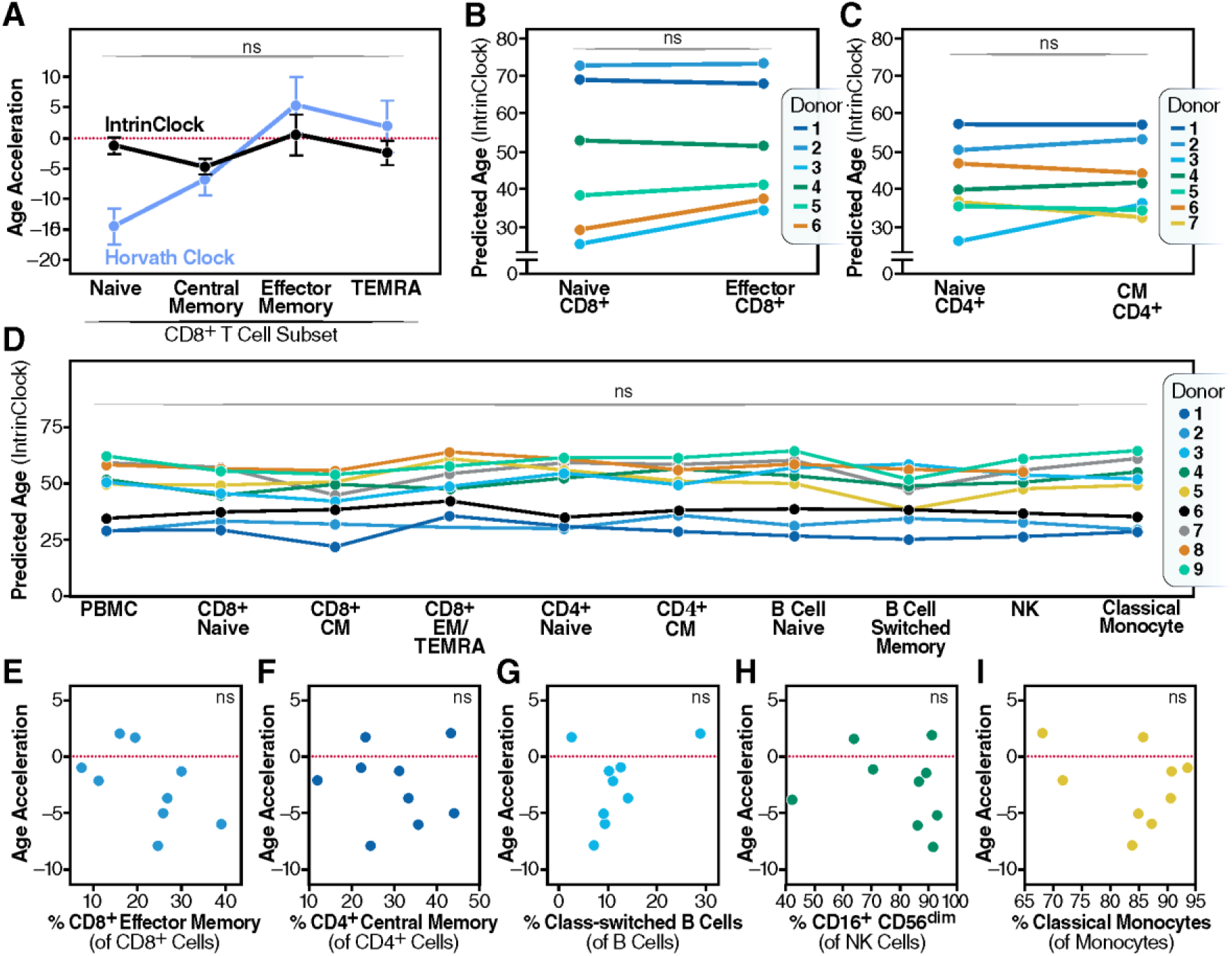
Epigenetic age accelerations by different clocks. **a**, Differences in epigenetic age accelerations in different CD8^+^ subsets generated in this study. Horvath clock predictions overlaid in light gray. **b**, Epigenetic ages of CD8^+^ naive cells and effector memory cells, based on data from GSE66564 and GSE83156. **c**, Epigenetic ages of CD4^+^ naive cells and central memory cells, based on data from GSE121192 and GSE71825. **d**, Epigenetic ages of PBMCs, CD8^+^ naive, CD8^+^ central memory, CD8^+^ combined effector and TEMRA, CD4^+^ naive, CD4^+^ central memory, B-cell naive, B-cell switched memory, CD16^+^CD56_dim_ NK, and classical monocyte cells. **e-i**, Association of percentage of e, effector memory CD8^+^ cells, f, central memory CD4^+^ cells, g, class-switched B cells, h, CD16^+^ CD56_dim_ NK cells, and i, classical monocytes with epigenetic age acceleration.

We also tested whether the IntrinClock would be similarly unperturbed in other immune cell types, particularly naive and memory B cells, which change in frequency with age^98^. We sorted CD8^+^ naive (CD8^+^CD28^+^CD45RO^-^), CD8^+^ CM (CD8^+^CD28^+^CD45RO^+^), CD8^+^ combined EM/TEMRA (CD8^+^CD28^-^), CD4^+^ naive (CD4^+^CD28^+^CD45RO^-^), CD4^+^ CM (CD4^+^CD28^+^CD45RO^+^), B-cell naïve (CD3^-^CD19^+^CD27^-^IgD^+^), class-switched B cells (CD3^-^CD19^+^CD27^+^IgD^-^), CD16^+^CD56_dim_ NK cells (CD3^-^CD19^-^CD56_dim_CD16^+^), classical monocytes (CD3^-^CD19^-^HLADR^+^CD14^+^CD16dim), and whole-peripheral blood mononuclear cell (PBMC) samples from nine donors aged 30–68 and collected DNA for methylation analysis. To increase cell recovery, we performed two sequential rounds of positive selection for CD8^+^ and then CD4^+^ cells using magnetic enrichment kits prior to flow sorting, similar to a published strategy^33^. Concurrently, we analyzed the PBMC samples using high-parameter spectral flow cytometry to empirically determine whether changes in immune cell composition of the PBMC samples would impact predicted epigenetic age of the whole PBMC fraction.

As predicted, we found no evidence for an association between cell subset and epigenetic age prediction (Figure 3D) or between cell subset and epigenetic age acceleration (ANOVA p-value > .05) (Figure S6A). This remained consistent whether epigenetic age acceleration was defined as the difference between predicted age and chronological age or as the residual after regressing predicted epigenetic age on chronological age. In contrast, cell subset and epigenetic age acceleration were significantly correlated, according to the Hannum (Figure S6B), Horvath (Figure S6C), Horvath Skin and Blood (Figure S6D), and PhenoAge (Figure S6E) clocks. To further investigate how resistant IntrinClock is to the change in immune cell composition, we analyzed the correlation between the PBMC epigenetic age and percentage of several PBMC subsets. As expected, we identified no significant relationship between the PBMC epigenetic age acceleration and percentage of CD8^+^ EM cells (Figure 3E), CD4^+^ CM cells (Figure 3F), class-switched B cells (Figure 3G), CD16^+^ CD56_dim_ NK cells (Figure 3H), or classical monocytes (Figure 3I), relative to their parent populations (Pearson’ s correlation p-value > .05). Combined with our observations of the IntrinClock’ s high accuracy across many tissues, these observations indicate that shifts in immune cell composition do not impact IntrinClock age predictions.

### IntrinClock is highly enriched for CpG sites upstream of transcription start sites, and its sites are enriched for motifs whose TFs are implicated in cancer

One central challenge in understanding epigenetic clocks comes from a lack of knowledge regarding to what extent epigenetic clocks are tracking a cell-autonomous or, conversely, a cell-ensemble phenomenon^99^. Our data provide evidence that current epigenetic clocks represent a composite of at least two variables, change in DNA methylation associated with aging in a cell intrinsic manner (IntrinClock), and a change in cell composition associated with aging. Due to the IntrinClock’s resistance to changes in immune cell composition, the CpG sites that constitute the clock may have more readily interpretable cell-autonomous biology as they are less likely to track markers of changing immune cell composition. This prediction could be particularly helpful in the context of identifying a functional or causal relationship between epigenetic clock sites and aging. We found that the sites in the IntrinClock that are hypermethylated with age are enriched within the region 200–1500 bp upstream of gene transcription start sites, and correspondingly strongly depleted in sites distant from genes (25% vs. 15%) (Figure 4A). In sites that are hypomethylated with age, there was a significant enrichment within the first exon of genes (8% vs. 5%) (Figure 4B). DNA methylation changes within 1500 bp of the transcription start site are most closely linked to alterations in gene expression^100^. Similarly, IntrinClock CpGs are enriched for being located near CpG islands (45% vs. 31%) and are depleted from open sea regions (20% vs. 36%) (Figure 4C).

**Figure 4.**
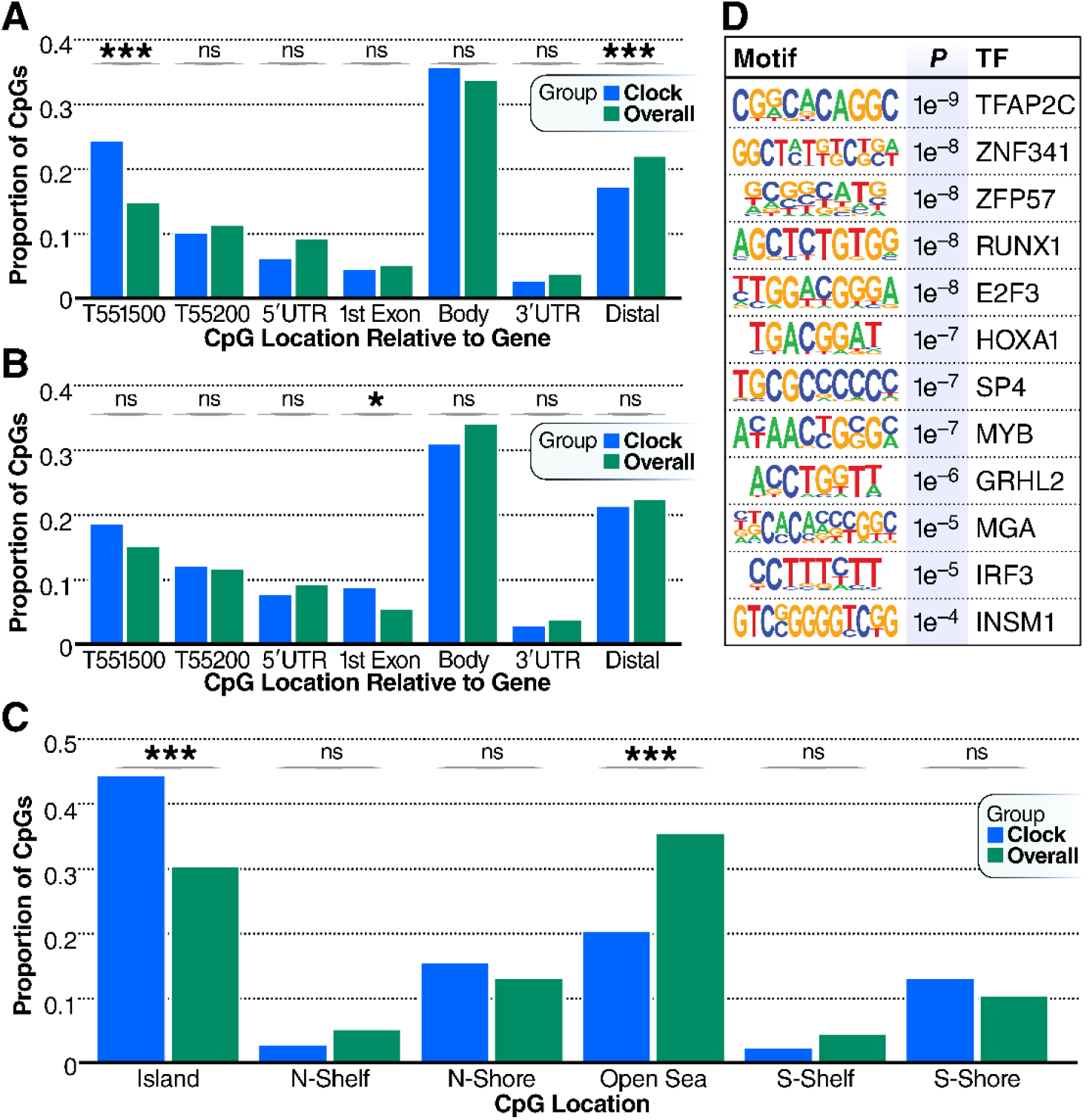
Distributions of CpG positions. **a**, Distributions of CpG positions relative to genes in IntrinClock sites that are hyper-methylated with age relative to background. **b**, Distributions of CpG positions relative to genes in IntrinClock sites that are hypo-methylated with age relative to background. **c**, Genomic distribution of IntrinClock CpG positions. **d**, HOMER analysis of the top 12 motifs enriched within 19bp on either side (5′ or 3′) of IntrinClock sites (40 bp total). *** one-sample proportion t-test p-value < .001; * < .05

Transcription factor activity and DNA methylation are biologically connected both directly, as in the case of E2F family transcription factors requiring methylated DNA to bind^101^, and indirectly, as in the case of passive methylation from lack of TF binding^102,103^. We investigated regions within 40 bp of IntrinClock CpG sites and used HOMER^104^ to identify enriched motifs associated with transcription factor-binding sites (Figure 4D). Motifs associated with TFAP2C, ZNF341, ZFP57, RUNX1, E2F3, HOXA1, SP4, MYB, GRHL2, MGA, IRF3, and INSM1 binding were significantly enriched, compared to a 40-bp background of basepairs surrounding CpG sites that are assayed by both Illumina Infinium HumanMethylation450K and MethylationEPIC chips. Aberrant activity of each corresponding transcription factor has been associated with cancer development or worsened prognosis^79,105–115^. Some of these, such as E2F3^116^ and IRF3^117^, have been associated with aging-related diseases, whereas a connection for others has yet to be discovered.

We were interested in exploring general patterns of shifts in IntrinClock CpGs with age. To avoid uneven distribution of tissue samples across age groups, we focused our analysis on blood samples. Given that a linear regression model was used to build the IntrinClock, we were not surprised that the two most prevalent patterns were a linear decrease and increase, respectively, of DNA methylation with age (Figure S7). However, we also found several CpGs (Clusters 4, 5, and 6) where the CpGs reverse their age-related direction of DNA methylation around the age of 21-30. This indicates that, for a subset of CpGs in the IntrinClock, there is a distinction between aging prior to and post sexual maturity.

Interestingly, these CpGs were 2.3-fold (34% vs. 14.9%) enriched for being located 200-1500bp upstream of a TSS, and 2-fold (19.4% vs. 10%) enriched for being located on a genomic south shore region (Figure S8), which are stronger enrichments than identified for IntrinClock sites generally (Figures 4A – 4C).

### IntrinClock epigenetic age is accelerated in models of intrinsic hallmarks of aging and in HIV^+^ individuals

HIV was one of the earliest conditions to be associated with acceleration of epigenetic age^74^. HIV infection is associated with a plethora of clinical manifestations and morbidities consistent with accelerated aging. However, HIV also causes major changes in immune cell composition^118^, which could skew previous versions of epigenetic clocks. As a result, it is unclear whether early results showcasing epigenetic age acceleration during HIV infection are due to changes in blood cell composition or an accelerated intrinsic rate of aging. Using the IntrinClock on previously generated data from HIV^+^ individuals and controls, we identified an HIV-associated increase in epigenetic age of two years, supporting the model that HIV leads to accelerated aging independently of shifts in immune cell composition (Figure 5A). We also sought to investigate whether the IntrinClock would be accelerated by other acute immune-related diseases. Using a dataset primarily generated in 2020, we found that the IntrinClock age prediction was not affected by COVID-19 (Figure 5B), contrary to findings in other epigenetic clocks where COVID-19 infection was associated with an increase in epigenetic age^119^. As the data analyzed in this study were generated early in the COVID-19 pandemic, most individuals would have been acutely, rather than chronically, ill with COVID-19. It remains to be seen whether the IntrinClock will predict a higher epigenetic age in those who are infected with COVID-19 for a prolonged period (i.e., long COVID).

**Figure 5.**
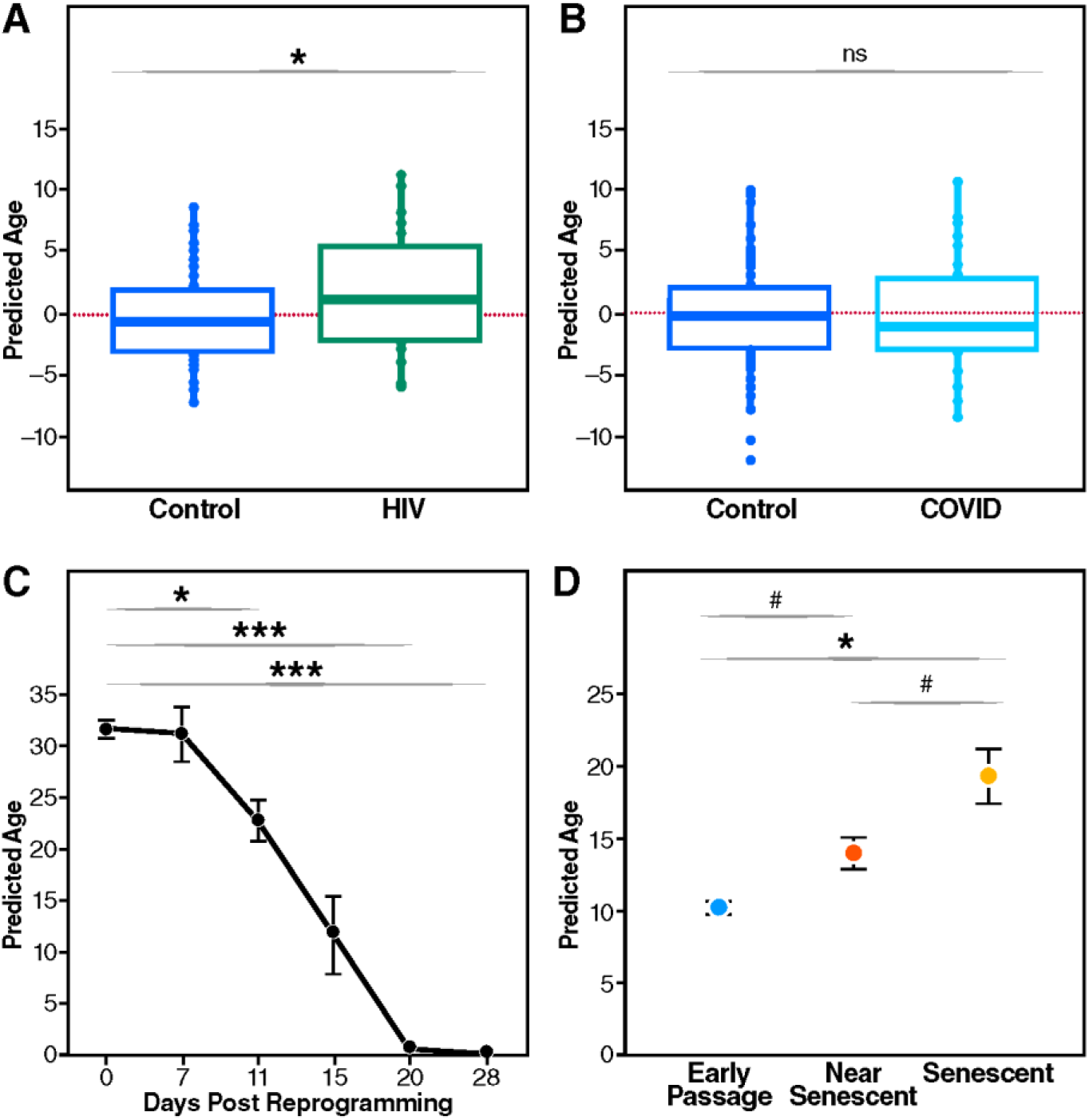
IntrinClock in HIV-infected individuals. **a**, Increase of IntrinClock epigenetic age in HIV^+^ individuals, DNA methylation data from GSE67751. **b**, No increase of IntrinClock epigenetic age due to COVID, DNA methylation data from GSE167202. **c**, Epigenetic reprogramming affects fibroblast predicted IntrinClock age. DNA methylation data from GSE54848. **d**, Induced replicative senescence in fibroblasts leads to an increase in IntrinClock predicted age. DNA methylation data from GSE91069. T-test p-values # < .10; * < .05; *** < .001.

One application of epigenetic clocks is in tracking the effect of rejuvenating or aging interventions on cells. As the IntrinClock was developed on sites that are not shifting due to immune cell compositional changes, we reasoned it may be more sensitive to such interventions. Consistent with this idea, we used an external dataset^120^ to find that the IntrinClock is sensitive to Yamanaka factor– mediated reprogramming in fibroblasts. The study authors sorted cells positive for TRA-1-60^+^, a marker for de-differentiation, at six time points after initiation of reprogramming. We investigated IntrinClock epigenetic age predictions at each time point and found that, from an initial mean predicted epigenetic age of 31, the age prediction decreased to 20 after 11 days of OSKM-mediated reprogramming. A mean age of 0 was reached after 20 total days of reprogramming (Figure 5C). Conversely, using publicly available data using an in vitro fibroblast model of replicative cellular senescence^121^, we found that the IntrinClock was progressively accelerated with cell divisions as cells become progressively more senescent. IntrinClock increased from a baseline predicted age of 10 to 15 after 14 population doublings, and then further increased to 20 after another 14 population doublings (Figure 5D). This effect was comparable to that seen using the PhenoAge clock, and stronger relative to the Hannum, Horvath, and Horvath Skin & Blood clocks (Figure S9).

## Discussion

Epigenetic clocks hold great promise for the study of longevity due to their high correlation with age and (particularly for second-generation clocks) association with aging-related disease state. As diagnostic tools, they have the potential to serve as important predictive biomarkers for assessing biological age, determining risk for age-associated diseases, and assessing the efficacy of interventions that target the aging process^122–126^. Recent technical advances, such as the development of principal component clocks^127^ and novel techniques for cost reduction^128^, promise to increase reliability and usability further. However, their current status as a composite of multiple aging signals makes them difficult to interpret and to link to specific biological processes. As an example, a recent study in patients post-COVID 19 infection demonstrated a significant PhenoAge epigenetic age acceleration in individuals over the age of 50, but an epigenetic age reversal for those under the age of 50^119^. Further, the manner in which clocks track healthspan is not fully overlapping, as clocks can be independently predictive of mortality even when analyzed jointly^129^. This challenge in interpretation is equally important for cellular models of the hallmarks of aging. In models of senescence or reprogramming, the sensitivity or even direction of the perturbation on predicted epigenetic age can dramatically differ, depending on the epigenetic clock used. For example, in this study, we identified the Hannum clock as predicting an age *reversal* in a fibroblast model of cellular replicative senescence (Figure S9).

The immune system changes dramatically with aging, and its decline can exacerbate or lead to many aging-related pathologies^130^. Clocks built solely on inflammatory markers can be used to predict age and risk of multimorbidity^131^. However, the presence of CpG sites that track primarily with immune cell markers makes epigenetic clocks applied to cell-intrinsic effects (e.g., cellular reprogramming in fibroblast cell culture) difficult to understand. Such sites can introduce background noise to the resulting measurement.

Here, using sorted CD8^+^ T-cell subsets, we observed that naive T cells consistently showed a younger epigenetic age than other CD8^+^ subsets (Figure 1C), ranging from a 10-year average age under-prediction in some clocks to as high as a 60-year underprediction in others. This suggests that epigenetic clock measurements are significantly affected by CD8^+^ T-cell differentiation. These observations reinforce the finding that current epigenetic clocks represent the integration of at least two variables: cell intrinsic aging and changes in immune composition during aging.

To isolate these variables, we developed a novel epigenetic clock that is based on CpG sites that do not change with CD8^+^ T-cell differentiation (IntrinClock). We further observed that this clock predicts the same age in each individual across a wide variety of immune cell types. Interestingly, a filtering step based on naive CD8^+^ T cells can generate a clock that is not affected by differentiation in cells from different lineages, such as CD4^+^ cells or even B cells. This indicates part of a unique “CD8^+^ naive” signal may, in fact, be a conserved quiescence program shared by a variety of immune cells. This observation is supported by our finding that methylation patterns associated with naive CD8^+^ T cells have a negative correlation with those changing with aging (Figure 2C). A connection between quiescence and aging is found in a wide variety of cell types, including neural stem cells^132^.

The IntrinClock’s higher proportion of sites near transcription start sites and CpG islands and its expected relationship with reprogramming and senescence suggest that it is tracking an intrinsic cellular aging program. Enrichment of IntrinClock CpG sites within motifs bound by transcription factors linked to cancer progression is consistent with a recent review investigating the connection between epigenetic clocks, global hypomethylation, cancer, and aging^133^. It will be important in the future to test whether acceleration of the IntrinClock is linked to particular disease states. This application could be a novel tool used to distinguish age-related diseases caused by aberrant cell-to-cell interactions from those caused by intrinsic cellular dysfunction.

The approach described here reduces the potential of cellular composition changes to be a confounder, particularly in blood or saliva samples, and will likely increase our understanding of biological aging and age-associated diseases. The IntrinClock holds the promise of being more sensitive to cell-intrinsic rejuvenation approaches, as its constituent CpG sites are not affected by immune cell composition. It may also be more closely linked to CpG sites with a functional or even causal relationship with the aging process. Overall, IntrinClock represents a new instrument to add to the aging biomarker toolkit, with a potential wide variety of applications and uses.

## Methods

### Ethics approval

NIH provided approval for use of phs000424/GRU (GTEx) age data via the dbGaP database approval system. Ethics approval was not required for other datasets generated.

### Immune cell isolation, sorting, and DNA extraction

PBMCs were extracted from leukopheresis chambers from CMV^+^ donors. Blood was first diluted 1:1 with PBS with 2% FBS. Diluted blood was slowly layered on top of 12 mL of Ficoll in a 50-mL Falcon conical tube. The tube was then centrifuged for 30 minutes at 2000 rpm at 21°C without applying a break. The layer containing white blood cells was removed, diluted with FBS-supplemented PBS, and centrifuged for 3 minutes at 2500 rpm. The cell pellet was re-suspended in 15 mL of ACK lysis buffer and incubated for 3 minutes. The cells were topped up with PBS with 2% FBS, centrifuged, and resuspended.

For the initial CD8^+^ epigenetic clock characterization experiment, an EasySep™ Human T Cell Isolation kit was used to extract T cells from the PBMC fraction. T cells were then washed, stained with 1:500 LIVE/DEAD™ Fixable Near-IR Dead Cell staining kit, washed, stained with an antibody cocktail (Supplementary Table 2), and washed again. FACS was performed on a BD FACSAria™ II instrument. DNA was isolated using a Zymo Quick-DNA™ Microprep Plus kit.

For the second comprehensive immune cell-sorting experiment, 2 million PBMCs were frozen immediately after extraction. The remaining cells were then positively selected for a CD4 fraction using the EasySep™ Human CD4 Positive Selection Kit II. The CD4 cells were stained with 1:500 LIVE/DEAD™ Fixable Near-IR Dead Cell staining kit, washed, and stained with CD4/CD8 antibody cocktail (Supplementary Table 2), and the remaining cells were positively selected for a CD8 fraction using the EasySep™ Human CD8 Positive Selection Kit II. Both CD8^+^ cells and remaining PBMCs were washed, stained with 1:500 LIVE/DEAD™ Fixable Near-IR Dead Cell staining kit and washed again. CD8^+^ cells were stained with a CD4/CD8 antibody cocktail (Supplementary Table 2), and the remaining PBMCs were stained with a B Cell/NK Cell/Monocyte antibody cocktail (Supplementary Table 3), after blocking with human IgG. All three fractions were then subjected to FACS analysis using a BD FACSAria™ II instrument. DNA was isolated using a Zymo Quick-DNA/RNA™ Microprep Plus kit.

For both experiments, DNA was quantified using Qubit™ HS dsDNA quantification reagents. Bisulfite conversion and DNA methylation assessment were performed by Diagenode. For all experiments involving FACS, post-sort validations were performed to verify cell sort purity by analyzing sorted populations via flow cytometry. The Clock Foundation assisted with facilitating DNA methylation assessment and data transfer for the initial CD8^+^ experiment.

### High-dimensional flow cytometry

PBMCs were transferred to a 96-well V-bottom plate. Cells were re-suspended in a 1:500 dilution of LIVE/DEAD™ Fixable Blue Dead Cell Stain kit in cold PBS and incubated for 30 minutes in the dark. Cells were then washed and blocked with human IgG for 30 minutes. They were then washed twice and stained with a PBMC phenotyping antibody cocktail (Supplementary Table 4). Cell phenotyping was performing on a Cytek Aurora™ instrument and analyzed using FlowJo™.

### DNA methylation analysis and pre-processing

.idat files were converted into beta values by using the *minfi* R package^134^, with a functional normalization pre-processing step^135^. For differential methylation analyses, beta values were converted to M-values through the formula M = log2(*B* / (1-*B*)). The R package *umap* was used for UMAP dimensionality reduction^136^.

### Dataset collection and pre-processing

All datasets used to build the novel epigenetic clock were either generated in this study or downloaded from GEO. Exact ages were obtained for GTEx data through dbGaP^137^, as exact chronological ages of tissues were required. For constructing the clock, the assembled database of DNA methylation data was first culled of any samples that had more than 10% of CpGs missing and of any CpGs that had more than 10% samples missing. All samples derived from cancer tissues were removed. To ensure forward compatibility, we filtered out CpGs that were not on the Infinium MethylationEPIC v2.0 array. Based on our CD8^+^ DNA methylation data, we tested the correlation of each CpG methylation with age and with naive CD8^+^ T cells. To assess whether CpGs were correlated with naive CD8^+^ cells, we binarized each naive sample as “ 1” and each non-naive (CM, EM, TEMRA) sample as “ 0” and then used the R *cor* function to compute a Spearman’s correlation between methylation and naive T-cell state. All CpGs with an absolute value correlation of .3 or greater with naive T-cell state were removed, and all CpGs with an absolute value correlation of .3 or less with age were removed.

Once CpGs and samples were filtered, the samples were split 75% for the training set and 25% for the test set. Imputation of missing was performed separately between training sets and test sets, and separately between different tissues within training sets and test sets (imputation performed using the *impute* R package^138^). Outliers were detected and removed using the *outlyx* function in the R *watermelon* package^139^. Untransformed beta values were used for model creation and age prediction. Prior to training the model, ages were transformed using Horvath’s formula used in his original epigenetic clock^2^. An elastic net model using *glmnet*^140^ was used to develop the IntrinClock, with alpha value set at .5. Once the first model was generated, the training data were a subset of only those CpGs with non-zero coefficients, which were used for training the final model.

### Statistical methods

For comparisons between two measurements from one individual, as in Figures 1B and 1C, paired t-tests were used for assessment of significant changes. For multiple comparisons between a group and a background reference, as in Figures 4A, 4B, and 4C, one-sample proportional tests using the *prop*.*test* function from the *R stats* package were utilized with Bonferroni multiple-comparisons correction. For samples of multiple measurements, repeated measures ANOVA implemented via the *statix* packages^141^ was used to test significance. Most graphs and figures were created with aid of the *ggplot* R package^142^.

### Motif enrichment and pattern analyses

For motif enrichment analysis, the HOMER software tool was utilized^104^. To define sequences of interest, we investigated 40-bp windows surrounding the 381 CpG sites that compose the IntrinClock. As a background, we investigated 40-bp windows around CpG sites in our dataset immediately before removal of CpG sites associated with naive CD8^+^ cells and those not associated with aging. For investigating patterns of IntrinClock CpG shifts with age, beta values from blood samples were converted to M values, after which the *degPatterns* function from the *DEGreport* R package^143^ was utilized. Patterns with fewer than 10 CpG sites were discarded from analysis. Ages were binned into groups of 10 (0-10, 11-20, 21-30, 31-40, 41-50, 51-60, 61-70, 71-80, 81^+^). Each bin was confirmed to have at least 100 samples.

### Epigenetic age acceleration analysis

To compute DNA methylation age for each epigenetic clock, the R *methylclock* package was utilized^144^. For experiments containing a limited number of donors or cell types, epigenetic age acceleration was defined as the difference between epigenetic age prediction and chronological age. For larger studies, epigenetic age acceleration was defined as the residual after regressing predicted epigenetic age on chronological age.

## Data availability

DNA methylation profiles generated in this study will be submitted to the GEO public repository prior to publication. Code used to generate the results in this study will be made public on Github prior to publication.

## Acknowledgments

This research was primarily funded by the Michael Antonov Foundation and Buck institutional support. Cloud computing support was provided by the Amazon Web Services Activate program, facilitated by the On Deck Longevity Biotech fellowship. Furthermore, we extend our warm appreciation to Dr. Steve Horvath and the Clock Foundation for assisting us with sample handling and data interpretation during the early phases of this work.

## Author contributions

N.A., H.M., and H.K. assisted in flow panel design and optimization. A.F. planned and conducted some of the experiments. R.R. and E.V. assisted in manuscript drafting. All authors contributed to manuscript editing and revising. A.T. designed, performed, and analyzed the experiments. A.T. drafted and wrote the manuscript.

