## Supplemental Figures and Tables for "Development of a novel epigenetic clock resistant to changes in immune cell composition"

### Supplemental Information

Supplementary Figures 1–3 describe the gating strategy used for FACS and for high-dimensional flow cytometry. Supplementary Figures 4–9 contain additional data. Supplementary Table 1 describes the datasets used for the training and test sets for model generation and validation. Supplementary Table 2 and 3 describe the antibody cocktail used for staining T cells and non-T cell PBMCs, respectively. Supplementary Table 4 describes the antibody cocktail utilized for high-parameter spectral flow cytometry.

### Supplemental Figures

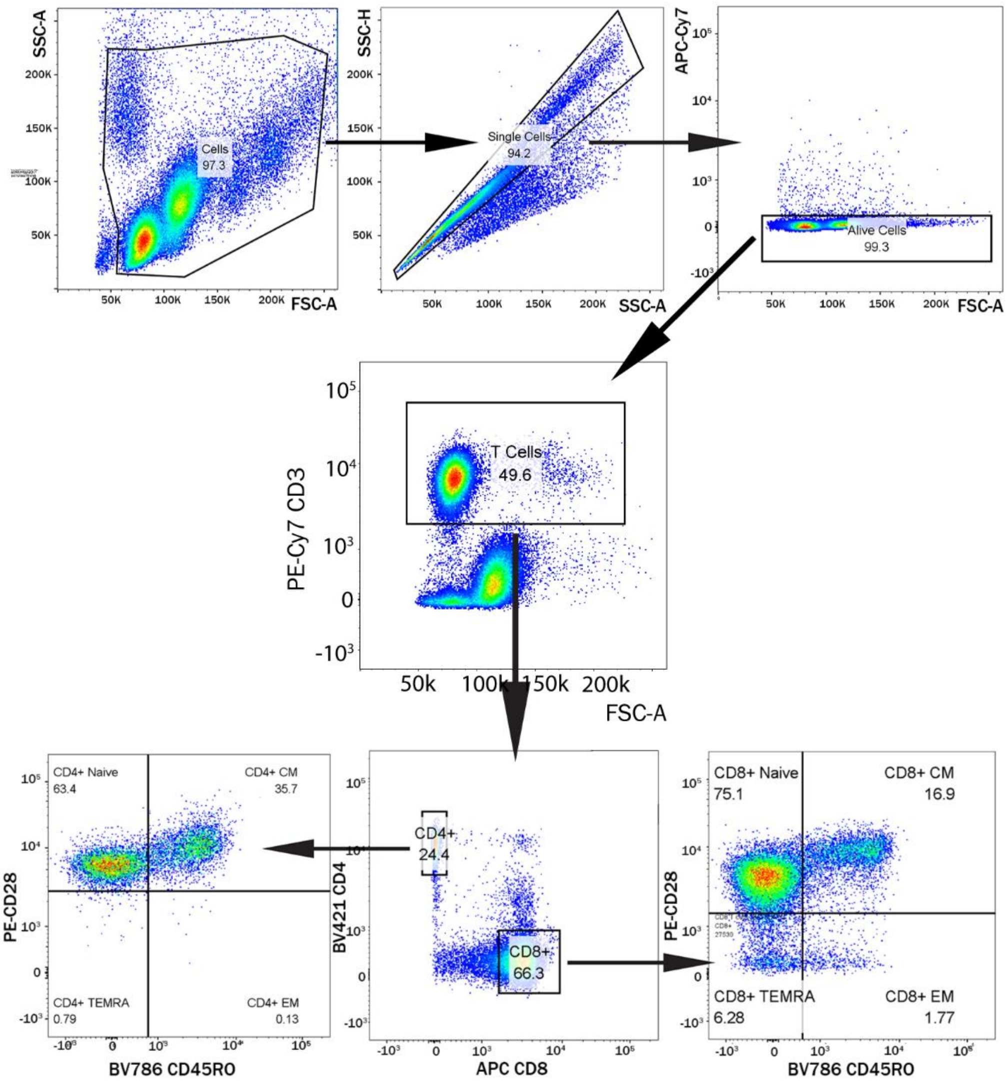

**Supplementary Figure 1. Gating strategy for sorting of CD4$^{+}$ subsets and CD8$^{+}$ subsets.**

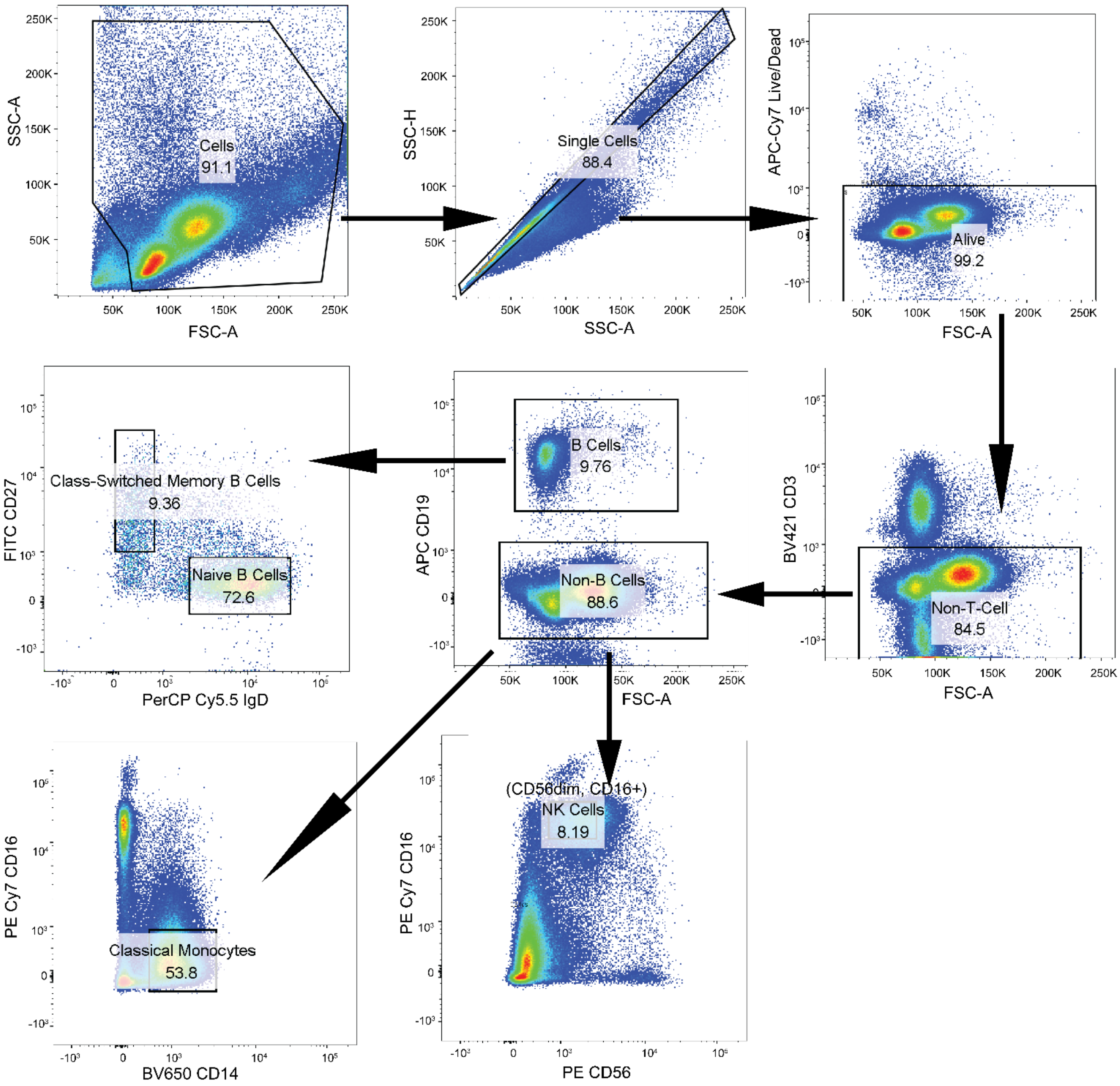

**Supplementary Figure 2. Gating strategy for sorting of B-cell subsets, $CD56_{dim}$ $CD16^{+}$ NK cells, and classical monocytes.**

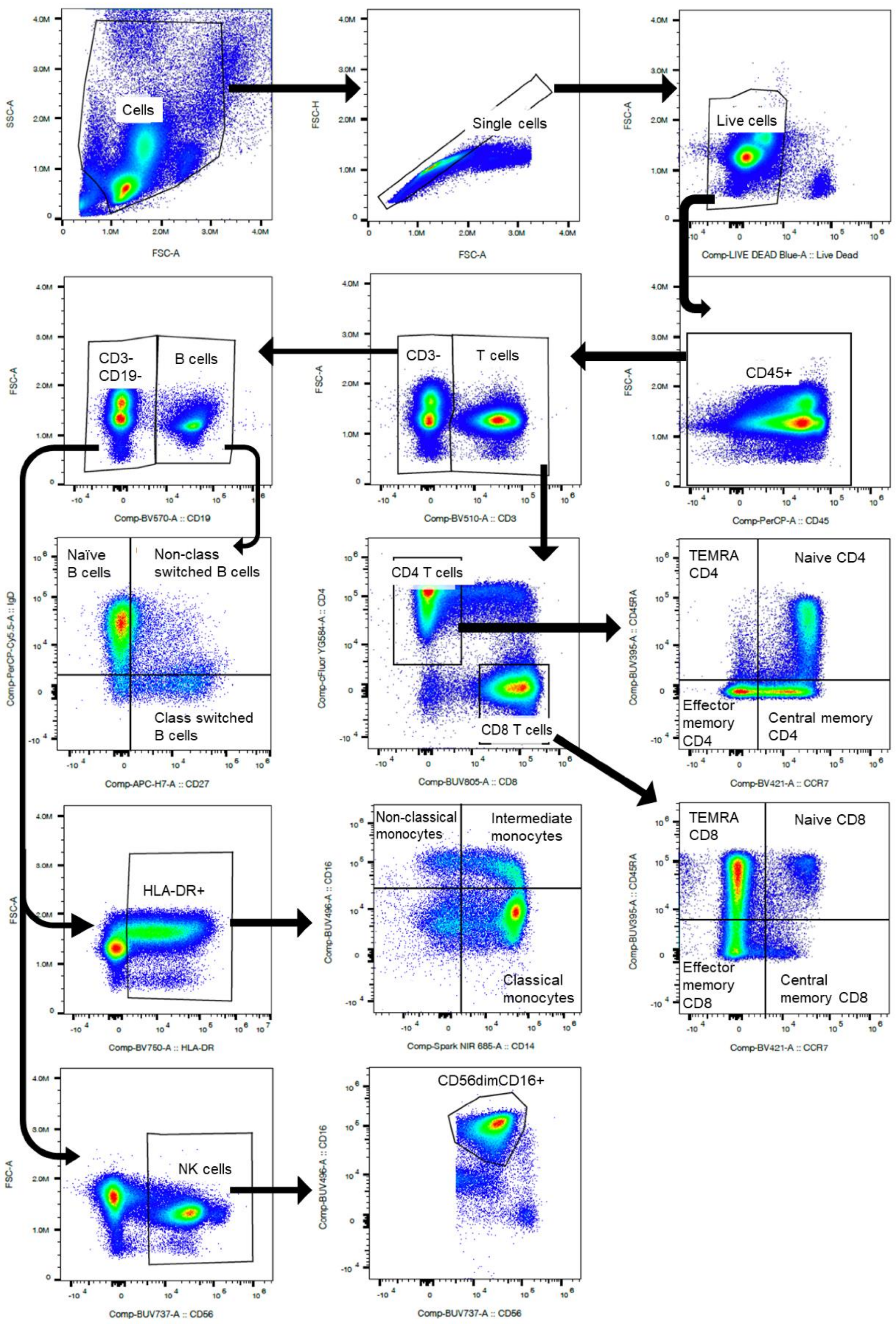

**Supplementary Figure 3. Gating strategy for high-dimensional flow cytometry, for data shown in Figure 3e-3i.**

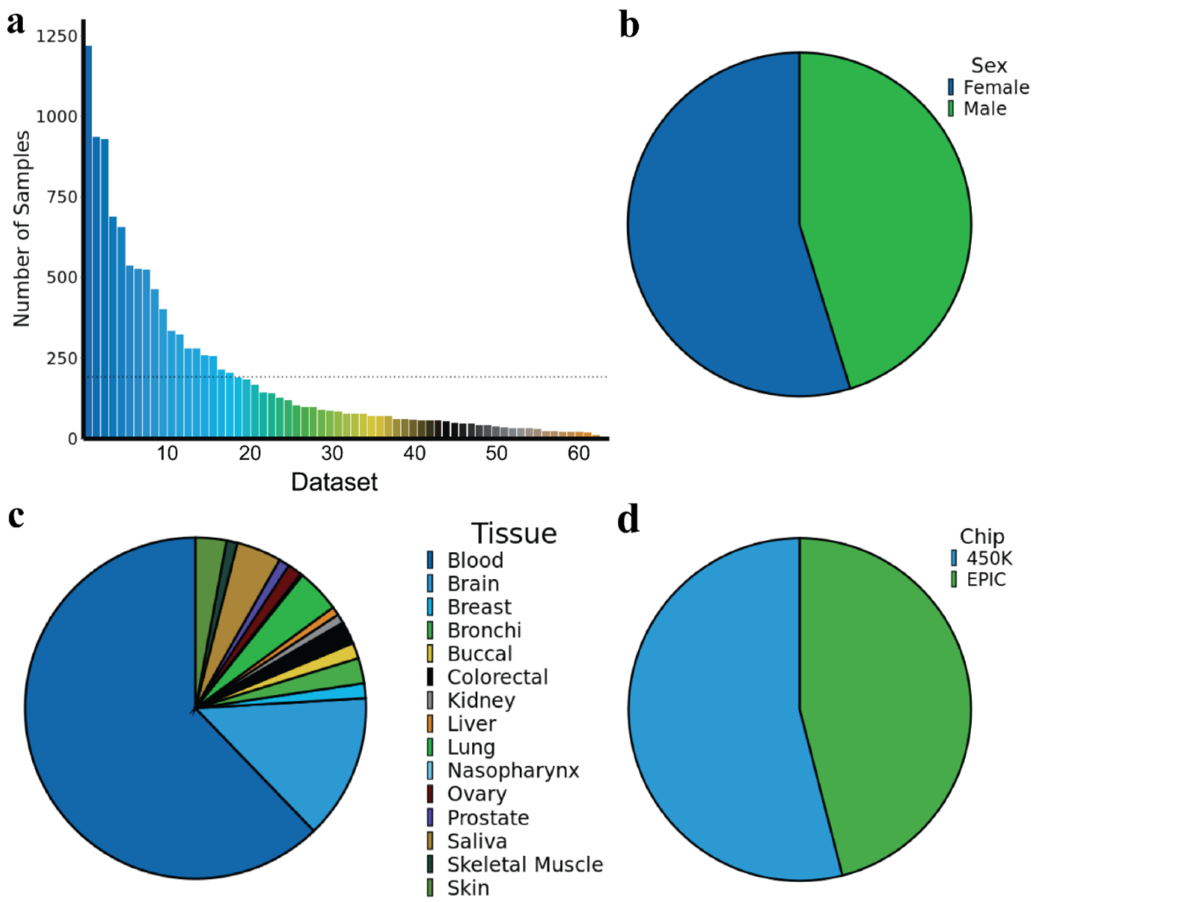

**Supplementary Figure 4. IntrinClock training and validation. a,** Number of samples used for IntrinClock training and validation per dataset. **b,** Sex distribution, **c,** tissue distribution, and **d,** Illumina chip distribution for samples used in IntrinClock training and validation.

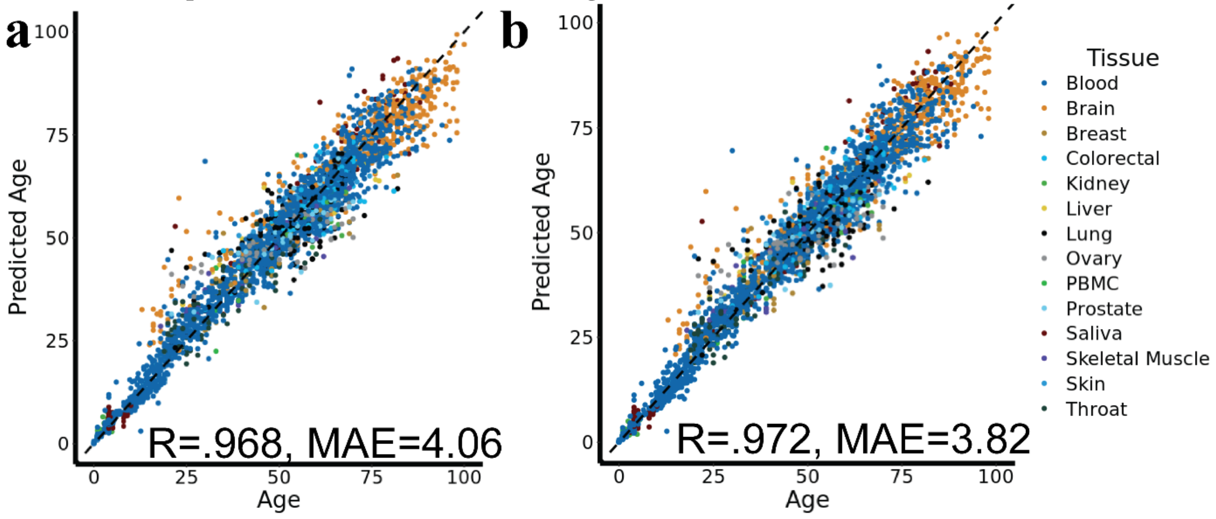

**Supplementary Figure 5. Prediction and accuracy of test sets. a,** Prediction accuracy and precision for the test set after training using one cycle of elastic net. **b,** Prediction accuracy and precision for the test set after training using once-repeated elastic net.

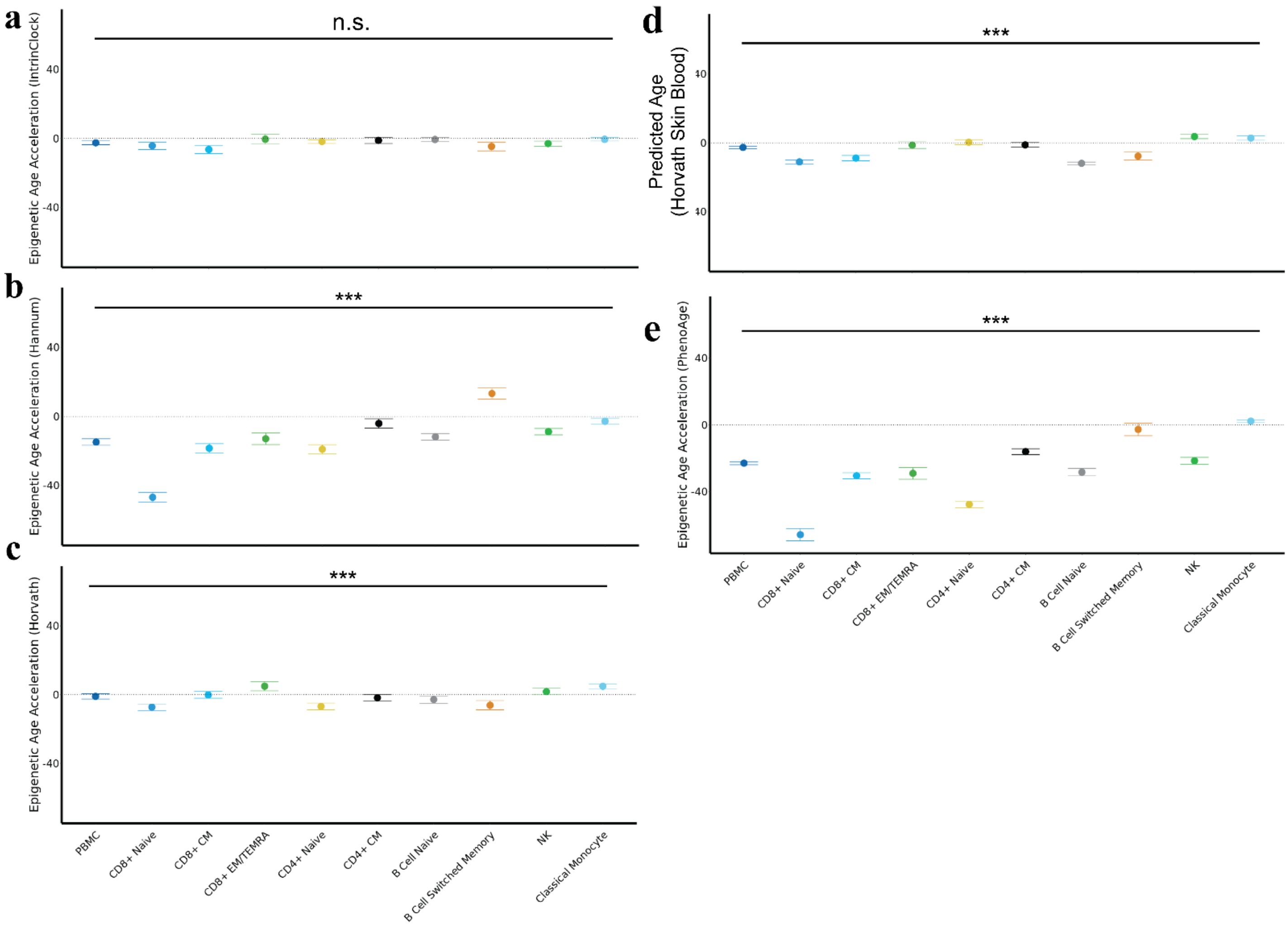

**Supplementary Figure 6**. **Predicted epigenetic age acceleration of matched-donor PBMC, CD8⁺ naive, CD8⁺ CM, CD8⁺ EM/TEMRA, CD4⁺ naive, CD4⁺ CM, B-cell naive, B-cell switched memory, NK, and classical monocyte samples by the a,** IntrinClock, **b,** Hannum**, c,** Horvath, **d,** Horvath Skin and Blood, and **e,** PhenoAge epigenetic clocks. *** ANOVA p-value less than .001. Error bars shown as mean ± standard error.

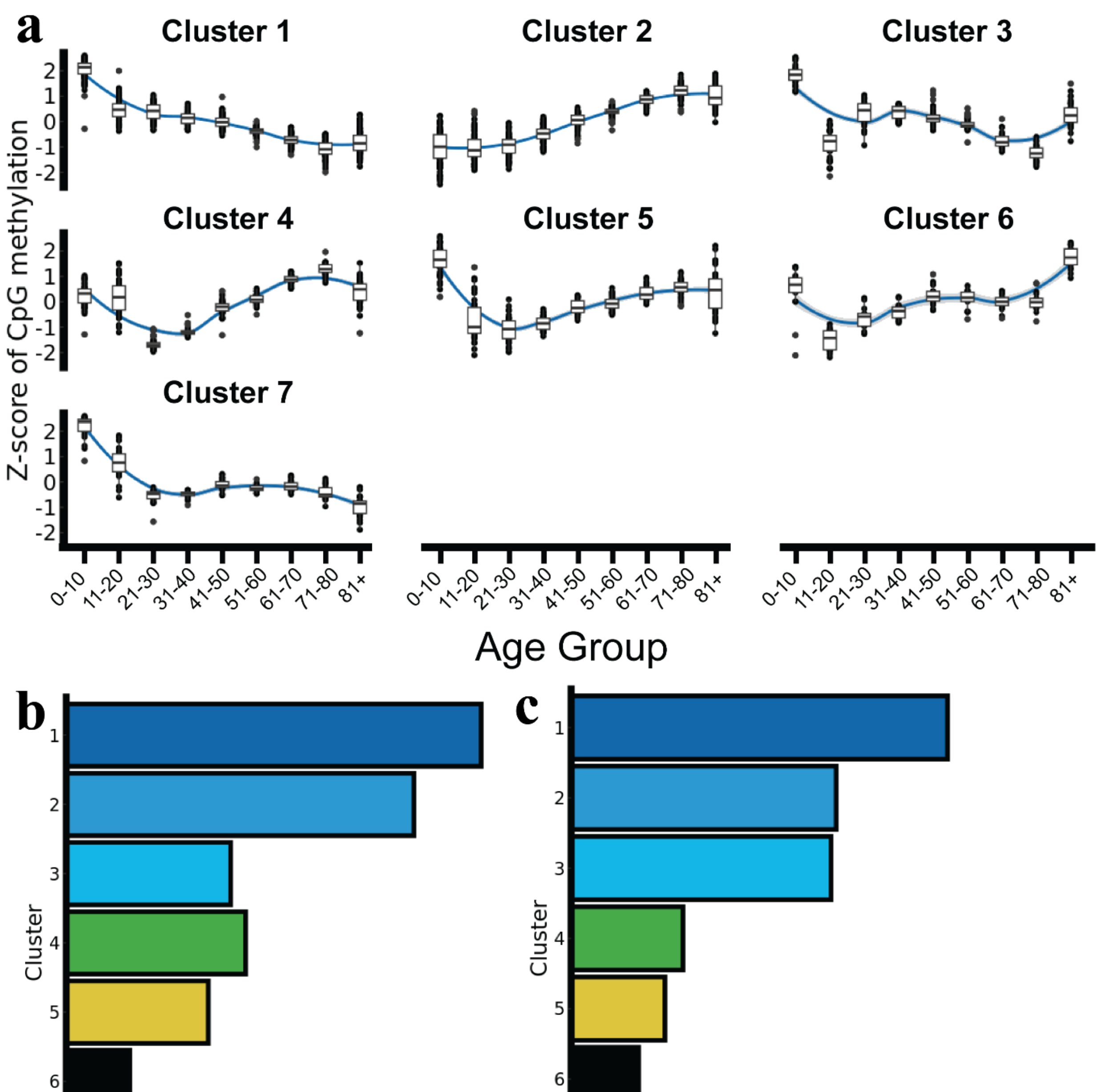

**Supplementary Figure 7. IntrinClock and CpG methylation patterns and clusters. a,** Analysis of IntrinClock CpG methylation patterns over five age groups in blood samples. **b,** Clusters by number of CpGs in the IntrinClock. **c,** Clusters weighed by the sum of absolute values of IntrinClock CpG coefficients.

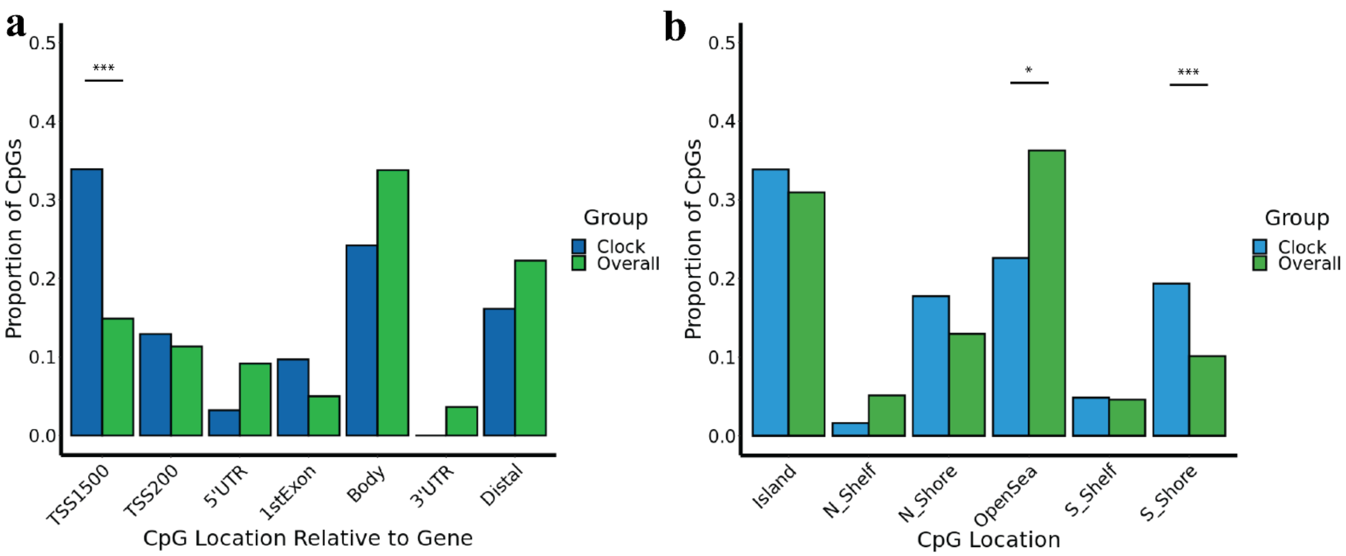

**Supplementary Figure 8. Analysis of IntrinClock CpG methylation sites belonging to clusters 4, 5, and 6 (identified in Supplementary Figure 7). a,** Distribution of clusters 4, 5, and 6 CpG sites relative to genes. **b,** Distribution of clusters 4, 5, and 6 CpG sites across the genome. *** One-sample proportion test p-value < .001; * one-sample proportion test p-value < .05.

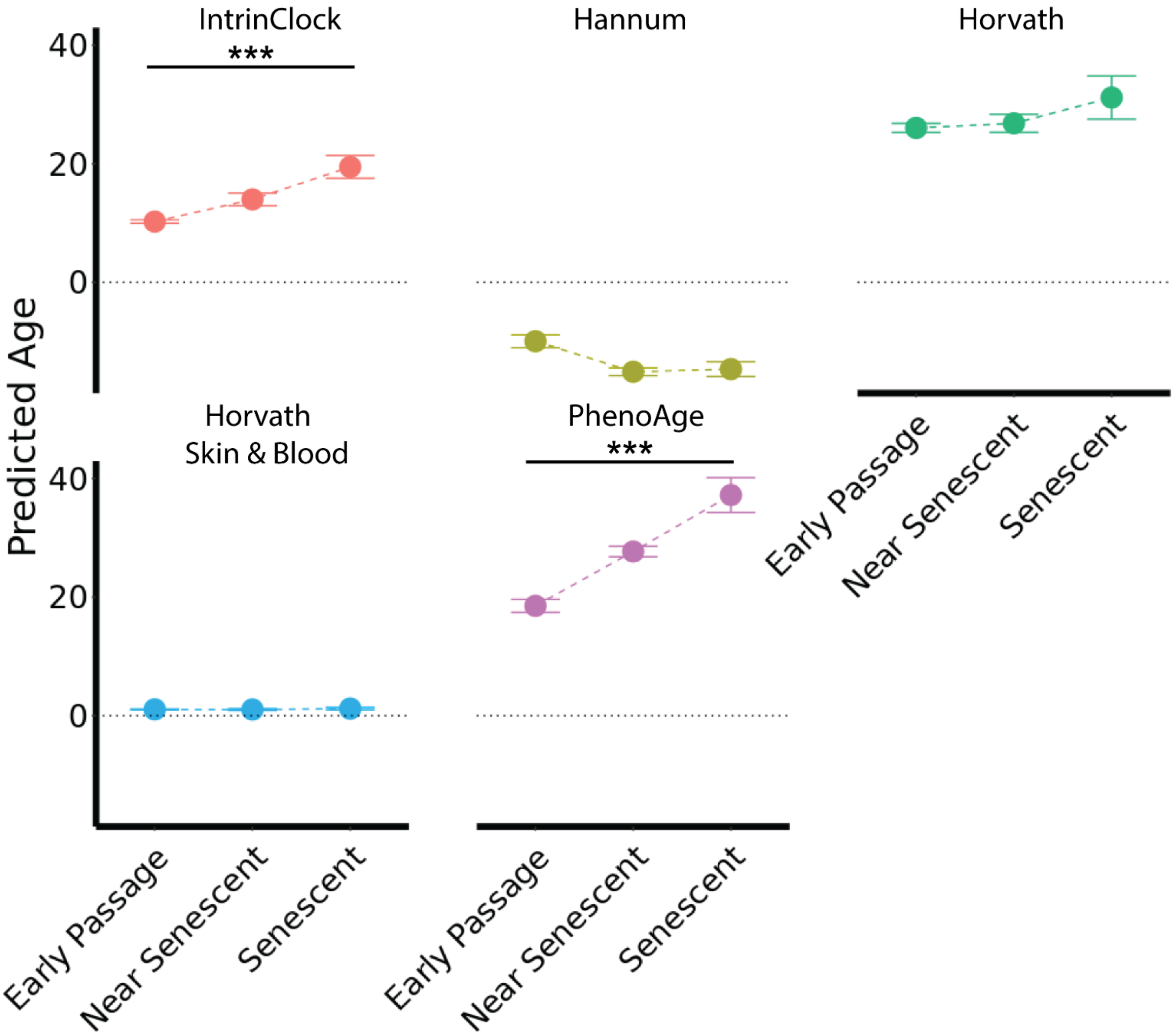

**Supplementary Figure 9. Effect of replicative senescence in fibroblasts on five epigenetic clocks.** DNA methylation data from GSE91069. *** t-test p-value > .001

**Supplemental Tables**

| **Supplementary Table 1. List of datasets used in this study for building and/or validating the IntrinClock.** | | | | |
|---|---|---|---|---|
| **GSE #** | Citation | Tissue(s) | N | Platform |
| **GSE41826** | Guintivano 2013 | Brain | 145 | 450K |
| **GSE42861** | Liu 2013 | Blood | 689 | 450K |
| **GSE42700** | Martino 2013 | Brain | 53 | 450K |
| **GSE115278** | Arpon 2019 | Blood | 474 | 450K, EPIC |
| **GSE52588** | Bacalini 2015 | Blood | 87 | 450K |
| **GSE201724** | Bartlett 2022 | Breast | 18 | EPIC |
| **GSE178887** | Bauer 2021 | Blood | 37 | 450K |
| **GSE191276** | Brennan 2022 | Blood | 6 | EPIC |
| **GSE136583** | Cerapio 2021 | Liver | 62 | EPIC |
| **GSE59157** | Charlton 2014 | Kidney | 92 | 450K |
| **GSE159898** | Li 2021 | Colorectal | 44 | EPIC |
| **GSE210245** | Clement 2022 | Blood | 36 | EPIC |
| **GSE112987** | Cobben 2019 | Blood | 103 | 450K |
| **GSE203399** | Cullell 2022 | Blood | 121 | 450K, EPIC |
| **GSE193879** | Davalos 2022 | Blood | 127 | EPIC |
| **GSE201752** | Estupiñán-Moreno 2022 | Blood | 113 | EPIC |
| **GSE129428** | Fries 2020 | Brain | 64 | EPIC |
| **GSE63347** | Horvath 2015 | Brain | 71 | 450K |
| **GSE179414** | Garcia-Prieto 2022 | Blood | 157 | EPIC |
| **GSE66351** | Gasparoni 2018 | Brain | 190 | 450K |
| **GSE99029** | Gopalan 2017 | Saliva | 57 | 450K |
| **GSE191200** | De Witte 2022 | Brain | 56 | EPIC |
| **GSE152026** | Hannon 2021 | Blood | 929 | EPIC |
| **GSE40279** | Hannum 2013 | Blood | 656 | 450K |
| **GSE119078** | Hearn 2020 | Saliva | 59 | 450K |
| **GSE92767** | Hong 2017 | Saliva | 54 | 450K |
| **GSE61256** | Horvath 2014 | Liver | 79 | 450K |
| **GSE78874** | Horvath 2016 | Saliva | 259 | 450K |
| **GSE141682** | Xiao 2021 | Blood | 42 | EPIC |
| **GSE120610** | McEwen 2018 | Blood | 156 | EPIC |
| **GSE40360** | Huynh 2013 | Brain | 46 | 450K |
| **GSE149282** | Ishak 2020 | Colon | 24 | EPIC |
| **GSE124366** | Islam 2019 | Buccal, Blood | 215 | 450K |
| **GSE52068** | Jiang 2015 | Nasopharynx | 48 | 450K |
| **GSE88883** | Johnson 2017 | Breast | 100 | 450K |
| **GSE69270** | Kananen 2016 | Blood | 184 | 450K |
| **GSE154566** | Kandaswamy 2021 | Buccal, Blood | 963 | 450K, |

| | | | | |
|---|---|---|---|---|
| | | | | EPIC |
| **GSE122288** | Kasuga 2022 | Blood | 61 | EPIC |
| **GSE157131** | Kho 2020 | Blood | 1218 | 450K, EPIC |
| **GSE167202** | Konigsberg 2021 | Blood | 525 | EPIC |
| **GSE70977** | Langevin 2015 | Buccal | 223 | 450K |
| **GSE141256** | Lewis 2020 | Small Intestine, Colon | 84 | 450K, EPIC |
| **GSE106648** | Kular 2018 | Blood | 279 | 450K |
| **GSE59685** | Lunnon 2014 | Brain | 526 | 450K |
| **GSE201872** | Magnaye 2022 | Bronchi | 142 | 450K, EPIC |
| **GSE114134** | Martino 2018 | Blood | 205 | EPIC |
| **GSE188593** | Muse 2022 | Skin | 64 | EPIC |
| **GSE85566** | Nicodemus-Johnson 2016 | Lung | 115 | 450K |
| **GSE166611** | Unpublished, Nonino 2016 | Blood | 32 | 450K |
| **GSE190540** | Vyas 2021 | Blood | 90 | EPIC |
| **GSE213478** | Oliva 2023 | Breast, Colon, Kidney, Lung, Skeletal Muscle, Ovary, Prostate, Testis, Blood | 987 | EPIC |
| **GSE112179** | Pai 2019 | Brain | 100 | 450K |
| **GSE203332** | Pihlstrøm 2022 | Brain | 336 | EPIC |
| **GSE137223** | Policicchio 2020 | Brain | 33 | 450K |
| **GSE61195** | Renauer 2015 | Blood | 21 | 450K |
| **GSE133062** | Ringh 2019 | Bronchi | 70 | EPIC |
| **GSE151017** | Ringh 2021 | Bronchi | 78 | EPIC |
| **GSE90124** | Roos 2017 | Skin | 322 | 450K |
| **GSE184269** | Roy 2021 | Blood | 167 | EPIC |
| **GSE112611** | Somineni 2019 | Blood | 402 | EPIC |
| **GSE178925** | Takeuchi 2022 | Blood | 24 | EPIC |
| **GSE146376** | Thompson 2020 | Lung | 280 | EPIC |
| **None yet** | This study | Blood | 30 | EPIC |
| **GSE50660** | Tsaprouni 2014 | Blood | 464 | 450K |
| **GSE72556** | Oelsner 2017 | Saliva | 93 | 450K |
| **GSE89707** | Viana 2016 | Brain | 49 | 450K |
| **GSE151407** | Voisin 2020 | Skeletal Muscle | 78 | EPIC |
| **GSE61107** | Wockner 2014 | Brain | 47 | 450K |
| **GSE49393** | Xu 2014 | Brain | 48 | 450K |
| **GSE174442** | Xu 2019 | Blood | 256 | 450K |
| **GSE128235** | Zannas 2019 | Blood | 537 | 450K |

| **Supplementary Table 2. Antibody mix used for sorting T cells.** | | | |
|---|---|---|---|
| **Target** | Fluorophore | Clone | Company |
| CD3 | PE-Cy7 | UCHT1 | BioLegend |
| CD8 | APC | SK1 | BioLegend |
| CD4 | Pacific Blue | OKT4 | BioLegend |
| CD28 | PE | B353546 | BioLegend |
| CD45RO | BV785 | UCHL1 | BioLegend |

| **Supplementary Table 3. Antibody mix used for sorting B cells, CD56$_{dim}$ CD16$^{+}$ NK cells, and classical monocytes.** | | | |
|---|---|---|---|
| **Target** | Fluorophore | Clone | Company |
| CD14 | BV650 | M5E2 | BioLegend |
| IgD | PerCP Cy5.5 | IA6-1 | BioLegend |
| CD3 | Pacific Blue | OKT4 | BioLegend |
| CD19 | APC | HIB19 | BioLegend |
| CD16 | PE-Cy7 | 3G8 | BioLegend |
| CD27 | FITC | O323 | BioLegend |

| **Supplementary Table 4. Antibody mix used for high-dimensional phenotyping of peripheral blood mononuclear cells.** | | | | | |
|---|---|---|---|---|---|
| **Specificity** | Fluorochrome | Vendor | Catalog # | Clone # | Lot # |
| **CD3** | BV510 | BioLegend | 344828 | SK7 | B364399 |
| **CD4** | cFluor YG584 | Cytek | R7-20042 | SK3 | F-012022-01 |
| **CD8** | BUV805 | BD Biosciences | 612889 | SK1 | 2006191 |
| **CD14** | Spark NIR 685 | BioLegend | 367150 | 63D3 | B343008 |
| **CD16** | BUV496 | BD Biosciences | 612944 | 3G8 | 1348178 |
| **CD19** | BV570 | BioLegend | 302236 | HIB19 | B350345 |
| **CD25** | PE-Fire 700 | BioLegend | 356146 | M-A251 | B357965 |
| **CD27** | APC-H7 | BD Biosciences | 560222 | M-T271 | 1260248 |
| **CD28** | PE-Cy5 | BioLegend | 302910 | CD28.2 | B336927 |
| **CD38** | APC-Fire 810 | BioLegend | 356644 | HIT2 | B365111 |
| **CD39** | BUV661 | BD Biosciences | 749967 | TU66 | 2075880 |
| **CD45** | PerCP | BioLegend | 386506 | 2D1 | B326914 |
| **CD45RA** | BUV395 | BD Biosciences | 740315 | 5H9 | 1356975 |
| **CD49d** | APC | BioLegend | 304308 | 9F10 | B270195 |

| | | | | | |
|---|---|---|---|---|---|
| **CD56 (NCAM1)** | BUV737 | BD Biosciences | 612766 | NCAM16.2 | 1210146 |
| **CD57 (HNK-1)** | eFluor 450 | Invitrogen | 48-0577-42 | TB01 | 2437632 |
| **CD95 (Fas)** | BV650 | BioLegend | 305642 | DX2 | B324443 |
| **CD127** | APC-R700 | BD Biosciences | 565185 | HIL-7R-M21 | 1341730 |
| **CD159a (NKG2a)** | Alexa Fluor 647 | BioLegend | 375105 | S19004C | B325094 |
| **CD161 (NK1.1)** | PE-eFluor 610 | Invitrogen | 61-1619-42 | HP-3G10 | 2446967 |
| **CD183 (CXCR3)** | PE-Cy7 | BioLegend | 353720 | G025H7 | B319137 |
| **CD184 (CXCR4)** | BV605 | BioLegend | 306522 | 12G5 | B301424 |
| **CD195 (CCR5)** | BUV563 | BD Biosciences | 741401 | 2D7/CCR5 | 1348905 |
| **CD196 (CCR6)** | BV711 | BioLegend | 353436 | G034E3 | B323435 |
| **CD197 (CCR7)** | BV421 | BioLegend | 353208 | G043H7 | B337639 |
| **CD279 (PD-1)** | BV785 | BioLegend | 329930 | EH12.2H7 | B351259 |
| **HLA-DR** | BV750 | BioLegend | 307672 | L243 | B368880 |
| **IgD** | PerCP-Cy5.5 | BioLegend | 348208 | IA6-2 | B366177 |
| **Klrg1** | PE-Fire 810 | BioLegend | 367733 | SA231A2 | B371259 |
| **TCRγδ** | BUV615 | BD Biosciences | 751308 | 11F2 | 2271540 |
| **Tigit** | BV480 | BD Biosciences | 747843 | 741182 | 2272802 |
